# SICaRiO: Short Indel Call filteRing with bOosting

**DOI:** 10.1101/601450

**Authors:** Md Shariful Islam Bhuyan, Itsik Pe’er, M. Sohel Rahman

## Abstract

Despite impressive improvement in the next-generation sequencing technology, reliable detection of indels is still a difficult endeavour. Recognition of true indels is of prime importance in many applications, such as, personalized health care, disease genomics, population genetics etc. Recently, advanced machine learning techniques have been successfully applied to classification problems with large-scale data. In this paper, we present SICaRiO, a gradient boosting classifier for reliable detection of true indels, trained with gold-standard dataset from genome-in-a-bottle (GIAB) consortium. Our filtering scheme significantly improves the performance of each variant calling pipeline used in GIAB and beyond. SICaRiO uses genomic features which can be computed from publicly available resources, hence, we can apply it on any indel callsets not having sequencing pipeline-specific information (e.g., read depth). This study also sheds lights on prior genomic contexts responsible for indel calling error made by sequencing platforms. We have compared prediction difficulty for three indel categories over different sequencing pipelines. We have also ranked genomic features according to their predictivity in determining false indel calls.

## Introduction

Reliable detection of short genomic insertion or deletion (Indel) still remains challenging for standard alignment-based variant calling methods (Wala et al. 2018). Indel detection has recently attained more focus from research community due to the advancement of next-generation sequencing (NGS) technologies. Short indels are genomic variants defined by insertion or deletion of one or more base pairs (defined as <50bp in (Alkan et al. 2011)) at a particular locus within the DNA. Although rarer than SNPs (Single Nucleotide Polymorphisms), they comprised 16% to 25% of all genetic variations, the second most abundant form of polymorphism (Mills et al. 2006; Mullaney et al. 2010).

Several genetic disorders are linked to deleterious indels such as cystic fibrosis, fragile X syndrome, trinucleotide repeat disorders, Mendelian disorders and Bloom syndrome (Kanehisa et al. 2015) and NGS has become the standard tool for disease variant discovery (Koboldt et al. 2013; Wang et al. 2013). Short indels are also assumed to cause some cancers, e.g., acute myeloid leukaemia, and lung cancer. Besides, indels can also modify promoter structures and affect gene expressions (Cheung and Spielman 2009). Indels are also used as genetic markers for populations (Vali et al. 2008). Due to their importance in population genetics and clinical genomics, accurate indel detection is of prime importance; more so in the case of germline de-novo and somatic mutations which are rare but mostly account for aberrant genomic alterations. NGS technologies (Quail et al. 2012) have continued to mature, become cheaper and are gradually getting adopted into clinical applications. Accuracy of SNP detection has reached at the level of 99^th^ percentile^1^. However, indel detection accuracy is not at the par, despite the advent of a large number of indel calling tools (Hasan et al. 2015). Highly used indel callers have an accuracy around 60% over whole genome (containing both high confidence and non-high confidence regions) (Cornish and Guda 2015; Hasan et al. 2015), which is in fact validated in this work. Performance studies over only high-confidence regions (regions for which we have a complete map of indels) report high precision. For example, on PrecisionFDA truth challenge^1^, DeepVariant (Poplin et al. 2018) reports a precision greater than 99% for indels only from high-confidence regions. When measured over whole genome, its precision drops to 63%. When we have taken concordance among multiple callers into account, performance drops even further. Indel calling error depends on sequencing platform characteristics, aligner and caller attributes as well as genomic context of the region (high vs. non-high confidence regions as mentioned in (Zook et al. 2016). Variant calling pipelines differ among their employed techniques and error landscapes. The techniques for one platform are hard to adapt across other platforms. The exact model of error distribution is also unknown (Poplin et al. 2018).

Over the years, lots of indel detection tools have appeared in the arena of NGS technologies. A comprehensive survey is out of the scope of this paper. Among the mainstreams, there are Samtools (Li 2011), GATK (McKenna et al. 2010a), Pindel (Ye et al. 2009), SOAPIndel (Li et al. 2013), VarScan (Koboldt et al. 2009), SplazerS (Emde et al. 2012), Dindel (Albers et al. 2011), SVM-M (Yang et al. 2016), IndelSeek (Au et al. 2017), Gindel (Chu et al. 2014), Sclapel (Fang et al. 2016), VarDict (Lai et al. 2016), LoFreq (Wilm et al. 2012) and more. A performance evaluation for some of them can be found in (Sandmann et al. 2017). All of them take aligned sequencing reads and make indel calls. To improve accuracy, ensemble of different methods has also been employed, e.g., HugeSeq [25] and BaySiC [26]. Data-centric approaches have also been explored, such as, SVM2 (Chiara et al. 2012), ForestSV (Michaelson and Sebat 2012), (Hwang et al. 2014) and Platypus (Rimmer et al. 2014). To address more difficult scenarios (e.g., de-novo and somatic indels) there exist tools like DeNovoGear (Ramu et al. 2013), SomaticSeq (Fang et al. 2015), and DNMFilter (Liu et al. 2014) among others. To validate these tools, projects have been launched to establish gold-standard datasets, e.g., Genome in a bottle (Zook et al. 2016, 2014, 2018) as well as SVclassify (Parikh et al. 2016). Recently deep learning techniques have made their way into genomic data analysis (Telenti et al. 2018; Zou et al. 2019), particularly in variant calling, for both short-read data (Illumina, Complete Genomics), e.g., DeepVariant (Poplin et al. 2018), GARFIELD-NGS (Ravasio et al. 2018) and long-read data (Pacific Bioscience, Oxford Nanopore), e.g., Clairvoyante (Luo et al. 2019).

In this paper, we have shown that, training of machine learning-based variant filters with the gold-standard datasets, using genomic contexts improves the performance of indel callers while retaining high sensitivity. Most, if not all, indel-callers (including above mentioned ones) rely on actual platform specific sequence reads which might be unavailable for large public datasets, e.g., exome aggregation consortium (ExAC) (Lek et al. 2016) and haplotype reference consortium (HRC) (McCarthy et al. 2015). Our filter can post-prune potential false indel calls to deliver a more reliable variant set.

Our specific contributions in this paper are summarized below:

1. We have developed a machine learning-based probabilistic filtering scheme to reliably identify false short indel calls. This improves state-of-the-art indel detection performance significantly, particularly when we consider whole-genome callsets (including non-high confidence regions or NHCR). We plan to make SICaRiO publicly available which can be trained and seamlessly integrated with any variant calling pipeline. For now, we will provide the tool and annotation data on request.
2. We have demonstrated that genomic contexts capture important prior information useful for reliable indel detection. To the best of our knowledge, no indel callers currently use this kind of features, e.g., GERP scores (Davydov et al. 2010), local DNA structures etc. We have also ranked genomic features according to their predictivity in determining false indel calls.
3. We have shown that different sequencing platforms make analogous calling mistakes albeit with different degree of similarity. This enables us to apply models trained with the callset generated from one platform over the callset generated from another platform. We have further estimated the performance for this cross-platform scenario.

## Results

### Performance Improvement After Variant Filtering

We have demonstrated the performance of our scheme by comparing the precisions before and after applying the variant filter. In ***Figure 1***, we have shown precision improvement of five different variant calling pipelines (each consisting a combination of sequenced base caller, read aligner and variant caller) over three different indel categories (see ***Supplementary Table 3***). The definitions of these three different indel categories are as follows:

1. Homopolymers: In this category, the inserted/deleted nucleotides as well as the previous or following nucleotide (the flanking prefix or suffix context) consist of only one type of nucleotides. Example: [A → AA], [AA → A]
2. Tandem repeats: The variant in this category is not a homopolymer; the number of inserted/deleted nucleotides is more than one and an inserted/deleted sequence is followed or preceded by the same sequence. Example: [AT → ATAT]
3. Aperiodic indels: Any other insertion/deletion which is not a homopolymer or a tandem repeat belongs to this category. Example: [A → AGT]

**Figure 1:**
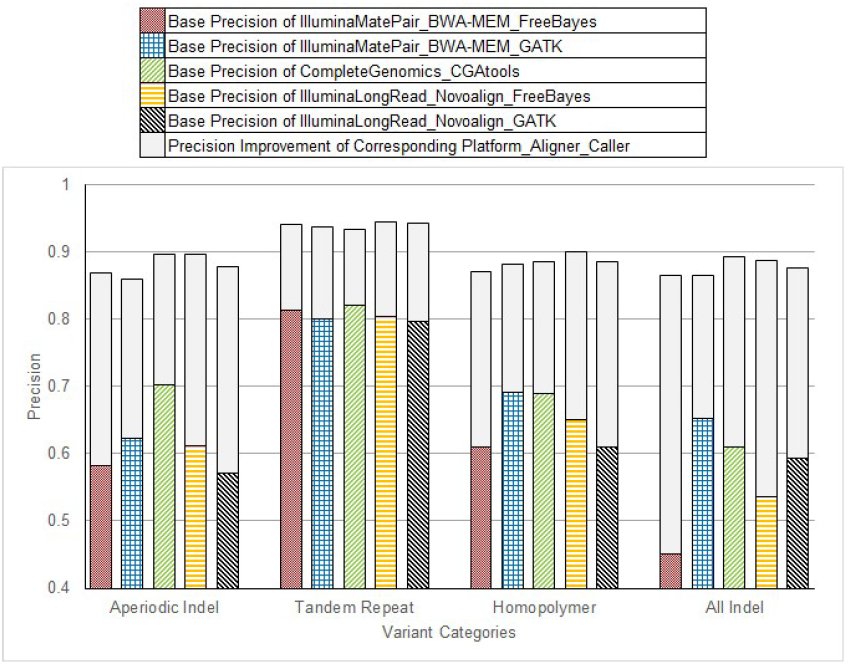
Precision Improvement across Five Different Platforms and Variant Categories.

We have also evaluated all indels together, when our training and test data contain all variant categories.

For aperiodic indels, before applying the filter, the maximum base precision of 70.3% is achieved by Complete Genomics (Peters et al. 2012). Illumina (mate pair^2^) with BWA-MEM (Li 2013) and GATK (McKenna et al. 2010b) ranks right after that with a base precision of around 62.22%. Illumina (long read^3^) with NovoAlign^4^ and GATK resides at the bottom of the table with a base precision of 57.16%. Application of our filter improves performances across all platforms/callers among which Illumina (long read) with NovoAlign and GATK shows the highest margin of improvement (30%). Complete Genomics shows the least albeit significant improvement (19%). The highest precision (89.70%) is achieved by Illumina (long read) with NovoAlign and FreeBayes (Garrison and Marth 2012) at a recall rate of 89.32%. The lowest precision (85.95%), not too far from the maximum, is achieved by Illumina (mate pair) with BWA-MEM and GATK at a recall rate of 88.44% (see ***Supplementary Table 3***).

For homopolymers and combined indel calls, most of the measures, i.e., base precision, filtered precision and platform recall show similar results like those of aperiodic indels. Here improvement is even bigger in some cases. For example, Illumina (mate pair) with BWA-MEM and GATK shows 41.5% precision improvement at a recall rate of 79.5% for combined indel calls. For tandem repeats, we have observed higher precision (93%) across all platforms at a superior recall rate (97%).

In ***Supplementary Table 4***, we have compared the precision improvement for combined indel callset of five more platforms including DeepVariant for which precision improvement is greater than 18.44% while keeping the platform recall at 86.25%. For details see ***Supplementary Table 4***.

We have investigated the effect of training sample size over precision to demonstrate the efficacy and extensibility of our filter in case of the availability of more samples having gold standard callset. We have used training sample size from one to four. All four training samples and the fixed test sample (HG002) are sequenced by Complete Genomics platform. The first data point in ***Figure 2*** [Left panel] is the base precision (no filter applied). This shows that the first training sample gives the largest improvement. Subsequently, the improvement slows down and after adding the fourth sample, tends to saturate. We have also investigated the effect using all gold-standard indel calls instead of only those true indels called by Complete Genomics as the true positive training callset. We can see both approaches produce similar curves but training with only Complete Genomics indel callset always performs better.

**Figure 2:**
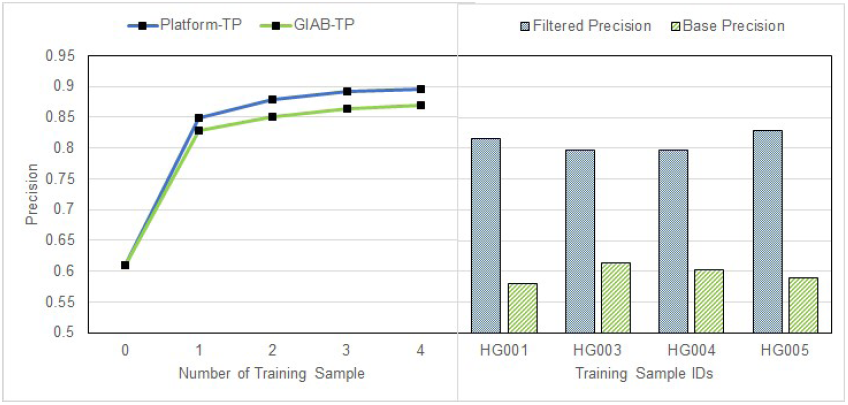
[*Left panel*] Precision vs. Training Sample Size for Complete Genomics Platform. True Positives Come from either True Platform Calls (Blue) or Genome-in-a-Bottle Gold Standard (Green). [*Right panel*] Filtered and Base Precision for Different Test Samples.

If training and test samples are different, there may exist a set containing indels common to both training and test callset. If the samples are from same population, this set will be larger and boost the classifier performance positively. To examine the interchangeability of the samples without common indel effect, we have used a single sample for both training and testing. From each sample we have trained with 80% indels and tested over the remaining non-overlapping 20% indels. We have used four unrelated samples, coming from three different population and observed small (3%) precision variability (see ***Figure 2***[Right panel]).

Although we have chosen precision as our performance measure (reason explained in *Methods* and *Discussion* section), the precision-recall curve of ***Figure 3*** shows that our filtering scheme has demonstrated high AUC or area under curve (0.965) for Complete Genomics platform. Since, AUC is a measure of ability to discriminate true and false indel calls, our filtering scheme can be used as an effective classifier. ***Supplementary Table 7*** corroborates this assumption as we can see good F1 scores (harmonic mean of precision and recall) over a range of threshold. We have used 0.5 as our threshold score which gives the highest F1 score (0.91). We can adjust the threshold according to our preference of precision and recall.

**Figure 3:**
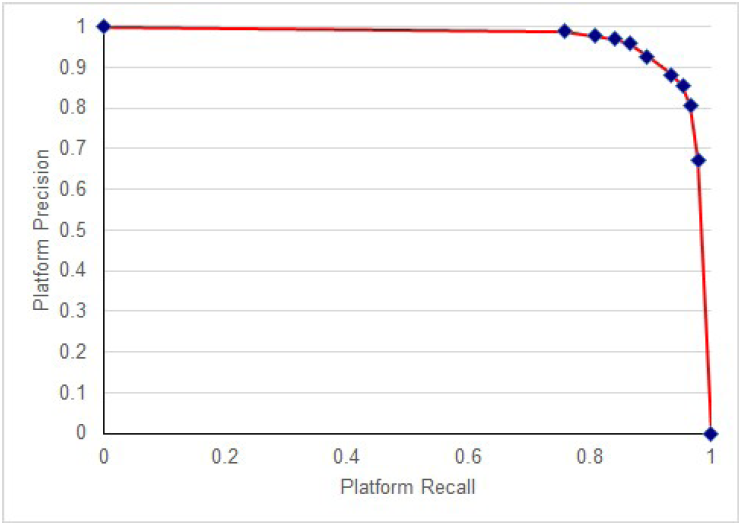
A Precision-Recall Curve for Complete Genomics Platform Trained over Sample HG001, HG003, HG004, HG005 and Tested over Sample HG002.

### Cross-platform Performance

Our platform-specific filtering scheme can be extended to the cross-platform scenario as well. We have trained nine different models from training callsets generated from nine different platforms (see ***Figure 4***). We have evaluated each of these trained models over nine different test callsets generated from the same nine platforms. The resulting matrix is shown as a heatmap in ***Figure 4***. Our first observation is that a model trained from a callset generated from a platform does not always perform the best on the test callset generated from the same platform.

**Figure 4:**
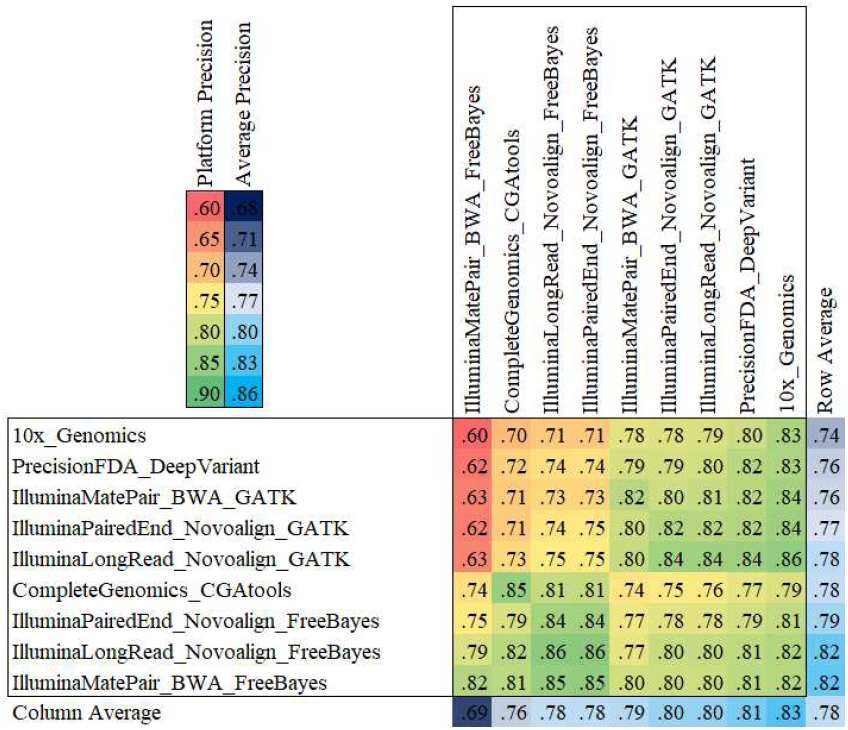
Cross Platform Precision Heatmap. Rows are Platforms Generating Training Indel Calls for Sample HG001/HG005 and Columns are Platforms Generating Test Indel Calls for Sample HG002.

For example, the model trained using callset generated by 10x_Genomics achieves 83% precision over the test callset generated by 10x_Genomics, whereas for the same test callset, model trained from data generated by Illumina (long read) with GATK achieves 86% precision. On the contrary, the best precision (85%) over Complete Genomics test callset is achieved by the model trained with Complete Genomics training callset.

Our second observation is that there are platforms whose generated callsets are hard to predict unless the model is trained with callsets generated from similar platform, e.g., Illumina (mate pair) with FreeBayes caller. On the other hand, there are platforms, whose generated callsets can be used to train models that can perform well over callsets produced from diverse platforms, e.g., Illumina (long read) with FreeBayes caller. So, these universal filters show the promise to be used in case we don’t have information about the data generating platform. An interesting observation is that if the train and test callsets share a common variant caller, e.g. FreeBayes, GATK, the corresponding model is likely to perform well. The greenish rectangle at the bottom-left corner of the matrix is created due to FreeBayes. Our cross-platform study demonstrates that using filter trained from any model will improve precision. Although some models will perform better than others, it is always better to use a filter than no filter.

### Feature Importance

We have grouped our selected genomic features used for classification into 10 feature groups according to their categorical similarities. XGBoost provides a score for each feature which indicates how useful the feature has been in the construction of the boosted decision trees within the model. By dividing each score with the maximum score, we have defined a measure named *relative importance*. For each group, we have calculated mean and standard deviation of the *relative importance* from the individual *relative importance* of member features (see ***Figure 5***).

**Figure 5:**
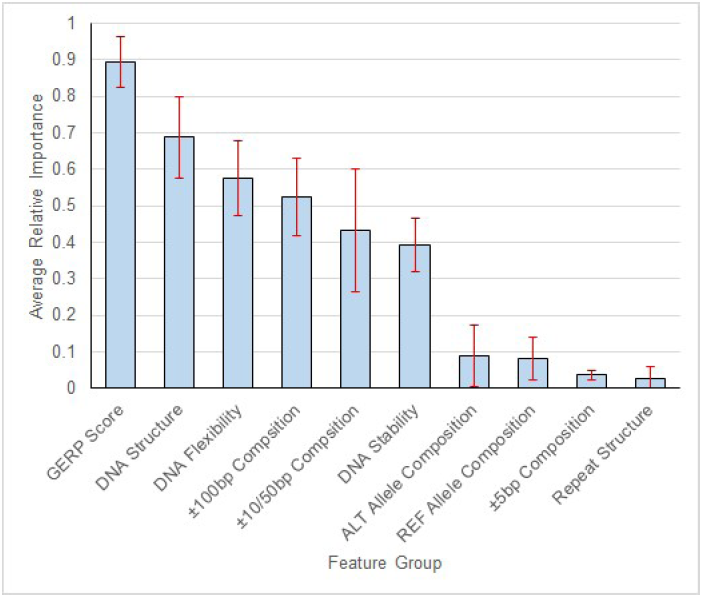
Mean and Standard Deviation (Red Error Bars) of Relative Feature Importance for Different Feature Groups.

We can see that the level of conservation (±10bp GERP score) is one of the best indicators of false indel calls. A feature is particularly useful in discriminating if its distribution differs between true and false positives. We have demonstrated that for 1bp downstream GERP score (see ***Figure 6***), true positives have a heavier-tailed distribution than false positives. Local physical DNA structure and flexibility are also good predictors. Longer genomic contexts, e.g., ±100bp distribution have better predictivity than shorter ones, e.g., ±5bp composition. An interesting observation is that the composition of reference or alternative allele has less importance than the composition of surrounding contexts in discrimination of true and false indel calls. A description of features is given in ***Supplementary Table 2***. and the ranking of features according to their importance is given in ***Supplementary Figure 4***.

**Figure 6:**
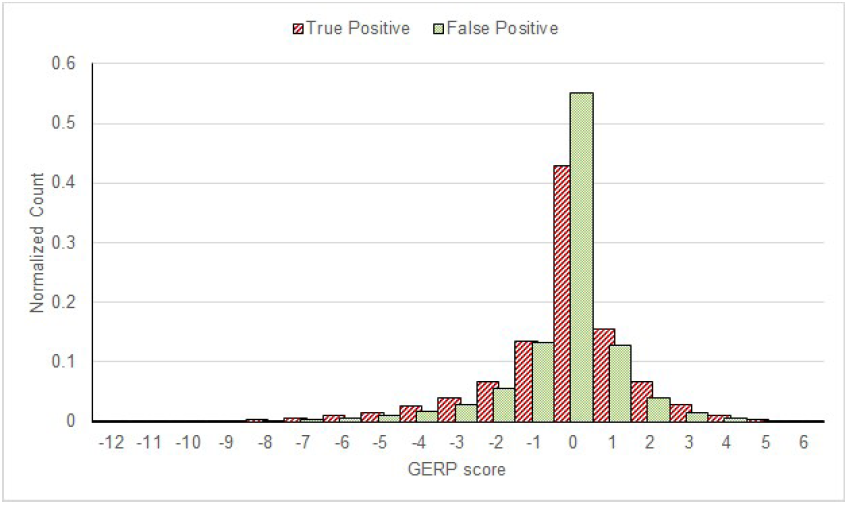
Histogram of 1bp Downstream GERP Scores of True Indel Calls (Red) and False Indel Calls (Green) from Complete Genomics Platform for Sample HG002.

## Discussion

Identification of true indel calls is of prime importance in many personal genomics applications. Due to the proliferation of next-generation sequencing, we have large repositories of indel calls at our disposals. With a few exceptions, these datasets do not come with detailed publicly available sequencing read information. To extract reliable indel calls from these large-scale data, our sequencing-read-independent filter should be advantageous. Our selected features can be computed from publicly available genome annotations.

Since, we are more interested in filtering false positives from called indels rather than identification of every rare indel call, we have reported precision as our primary performance measurement. However, our training algorithm minimize a loss function for binary logistic regression which does not prioritize precision in any way. We have provided all standard classification performance measures, i.e., recall, F1-score and area under precision-recall curve where appropriate. We have demonstrated that our filtering scheme has done well in all criteria; our choice of threshold has achieved the highest F1-score (.91) with an AUC > 0.965 for Complete Genomics platform. Our reported recall estimates the fraction of platform-supported true indels (only those called by platform) passed by our filter as positive; this does not estimate the fraction of all GIAB-supported true indels (the gold standard, some of which are not called by the platform) passed by our filter as positives. This makes sense, as our filter will not get a chance to evaluate an indel not called by the platform. Thus, we cannot know the true recall of our filter, and this is another reason to use precision as our metric.

We have demonstrated the efficacy of GERP scores in recognition of true indel calls. ***Figure 6*** shows a histogram of 1bp downstream GERP score for both true and false indel calls. Indel calls around the region with neutral GERP scores exhibit a strong bias for false indel calls. One possible explanation of this phenomenon is that regions under no selective pressure can easily accumulate repetitive elements (e.g., homopolymer, tandem repeats). Sequencing of these regions are inherently difficult and error-prone.

Our training and validation data consist of gold-standard indel calls from both high and non-high confidence region (NHCR). Since we do not have a complete map of NHCR indels for gold-standard samples, most of the performance evaluation tasks reported in the literature have been conducted using only indels from high confidence regions (e.g., PrecisionFDA truth challenge^5^). In our opinion, this approach has two drawbacks. First, density of false indels calls are much higher (10% of the genome harbours greater than 50% of false indel calls) in NHCR, despite the fact that a fraction of these NHCR indel calls (in our estimate around 3%, see ***Supplementary Table 9***) is expected to be true. Second, for a new sample we do not know exactly which portions of the genome are high confidence regions. Thus, discarding NHCR indel calls beforehand is not a feasible option. We argue that the inclusion of NHCR indel calls in our training set improves our model’s capability of false indel call detection without putting us in any advantageous situation with respect to precision, our principal performance metric. Rather our reported precision can be seen as an underestimation of true precision (which could be calculated if we would know the complete NHCR indel map), since our training set is more biased towards type II error (from the viewpoint of statistical hypothesis testing) or false negative (missing true indel calls).

We have observed that the performance of our classifier is consistently better in the context of different train-test sample than in the context of same sample train-test splits. We hypothesize that in case of a different train-test samples, a set of indel calls are common in both samples. In actual scenario, we are most likely to apply the classifier across samples and get that additional performance boost.

One of the current tides in genomic data analysis and in the broader area of machine learning is the extensive application of deep neural network architecture. We have previously mentioned DeepVariant which shows promising results for variant calling. The biggest downside of deep architectures is the difficulty of model interpretation within the black-box, although researchers are actively working on this issue (Shrikumar et al. 2017). Unlike other areas, interpretability is very important in biology, as this lay the foundation of the broader understanding about living systems. Another downside is the existence of adversarial examples, a sample with marginal noise which will be disproportionately misclassified by the model. Requirement of large training data (with no clearly reliable techniques for genomic data augmentation), high-end GPU platform, complicated hyperparameter tuning add up to the complexity. In our context, our use of gradient tree boosting directly gives feature importance. Moreover, our annotation data is mostly static (as we don’t use read data and model particular error pattern) and will not vary with more samples. Rather, more training data will saturate our understanding about unreliable genomic contexts corresponding to a particular sequencing platform.

A potential limitation of our filter is that it may have less sensitivity towards rare or de-novo variants. Discrimination between a sequencing error and a rare variant is an inherently complicated task and our filter also suffers in this context, more so since we don’t use sequencing reads. However, at a bare minimum, it can provide a prior probability of being a true indel to be used with other evidences for a hard to validate indel.

## Methods

We have formalized the identification of false indel calls as a supervised binary classification problem. Our basic assumption is that the genomic context of a true indel callset and that of a false indel callset differs from each other. Decision tree is an effective model for classification problems. Ensembles of decision trees are some of the most powerful off-the-shelf classifiers available to data analysts (Friedman et al. 2000). We have particularly used XGBoost (Chen and Guestrin 2016), an implementation of decision tree-based gradient boosting classifier. XGBoost provides a verdict on being a true indel as well as an estimate of uncertainty with a probability score. For training and testing, we have used genome in a bottle (GIAB) data set. To train a binary classifier we need instances from both classes, i.e., true and false indel calls, annotated with relevant genomic contexts captured within a feature matrix. Our positive instances have come from the indel callset generated by the corresponding platform and supported by GIAB high-quality gold standard dataset. The generated indel callset not present in the gold standard dataset are our negative instances for corresponding platform. We have annotated both positive and negative instances using publicly available genomic features, not related to any specific platform or caller or population.

### Dataset

We have used both platform-specific indel calls and the gold-standard indel calls from GIAB (Genome-in-a-bottle) consortium. In our primary evaluation, for training purposes we have used indel callsets of three samples, i.e., HG003, HG004 and HG005, generated from five different sequencing pipelines (see ***Figure 1***). For some platforms, indel callset of only one training sample (HG001) is available which we have used to evaluate them (see ***Supplementary Table 4***). For testing purposes, we have always used sample HG002, as this was sequenced by each platform. To assess the effect of the sample size over performance, for training purposes we have added indel callsets from four samples generated from Complete Genomics, namely, HG005, HG004, HG003 and HG001, in the given order. For annotation purposes, we have used RepeatMasker (Tempel 2012) for identifying various repeat regions, e.g., tandem, interspersed etc.. To measure the level of conservation, we have used GERP scores (Davydov et al. 2010). All DNA sequence contexts are calculated with hg19 reference genome (Rosenbloom et al. 2015).

### Feature Selection

Extracting representative features is a crucial step for building any machine learning classifier. Here the goal is to select features which are relevant predictors for sample class, e.g., true vs false calls. We have tried a number of genome annotations as well as their combinations to find a good set of platform-independent genomic features. We have not used any platform or population specific features, such as, read depth, quality, alignment, allele frequency, genotype calls, etc. All of the features can be generated with publicly available genome sequences and annotations.

We have found that functional features, e.g., gene annotation from GENCODE (Harrow et al. 2012), open chromatin, binding sites, histone modification etc. from ENCODE (Bernstein et al. 2012) are less indicative of indel calling accuracy; instead physical properties (Thomas Abeel 2011), nucleotide composition (distribution of monomer, dimer and trimer), level of conservation (GERP scores) (Davydov et al. 2010) and repeat structure of the indel and its surrounding DNA are more suggestive. We have calculated these features for four corresponding contexts of different scales, namely, ±5bp, ±10bp, ±50bp and ±100bp. In total, we have kept 219 features which are found to discriminate true indels from false ones.

### The Model

Boosting is an ensemble technique in machine learning that combines several weak learners to build a strong learner (Friedman et al. 2000). We can formulate boosting using a general class of mathematical model, i.e., the adaptive basis function model. Here, for a given dataset *x*, a target function *f(****x***) is expressed as a linear combination of *d* basis functions as in the following equation:

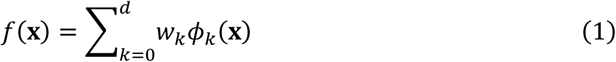

Here, ϕ _*k*_(***x***) is the *k*^th^ basis function, i.e., weak learner/classifier, and *w*_*k*_ is the assigned weight of *ϕ* _*k*_(***x***) in decision making. In gradient tree boosting, each *ϕ* _*k*_(***x***) is a decision tree classifier.

Behind most supervised learning algorithm, the assumption is that optimal model parameters will minimize the risk of misclassification. True risk of misclassification is approximated through the introduction of some specified per-sample loss function ℓ *(y*_*i*_, *f(****x***_*i*_)) for the *i*^th^ sample, where *y*_*i*_ is the true class label. A general stage-wise boosting algorithm defines a recursive relation between the ensemble learner from stage *k* and *k – 1* for the *i*^th^ sample as follows:

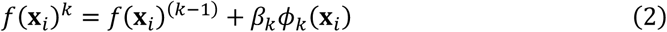

Learning *(β* _*k*_, *ϕ* _*k*_) can be cast as an optimization problem which tries to minimize the risk according to following equation:

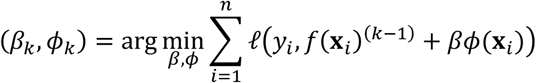

Gradient boosting splits this single step optimization into two parts. First it fits *ϕ*_*k*_*(x*_*i*_) to a new quantity called pseudo-residuals *r*_*ik*_ where,

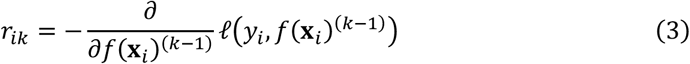

In fact, *r*_*ik*_ is the negative of the gradient of loss function for the *i*^th^ sample with respect to *f(x*_*i*_)^*(k–1)*^ from stage *k – 1*. This is an example of functional gradient descent giving the direction of steepest descent in the space of loss function. The amount of total descent or the weight, *β*_*k*_ of the basis function *ϕ*_*k*_*(x*_*i*_) is calculated using the following equation:

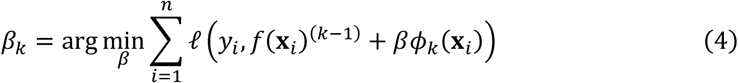

Sometimes, instead of Equation 1 we use the following recursive equation:

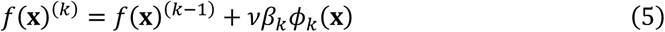

Here, *v* is called a shrinkage factor which is used to slow down stage wise model building to avoid overfitting.

In our experiments, we have used XGBoost (Chen and Guestrin 2016), an efficient, scalable, sparsity-aware, cache-oblivious, distributed implementation of gradient tree boosting. It also uses second-order Taylor series approximation of a regularized loss function for fast computation.

Decision tree-based learning models provide a natural framework for feature importance estimation. Gradient tree boosting extends the framework for an ensemble of decision trees. Since the importance of each feature is calculated separately without being entangled with other features, we can sort the relative importance of features and provide a global ranking. At every non-leaf node of a decision tree, a feature with an appropriate value (splitting point) is selected which will reduce the node impurity (mixture of samples from different classes) maximally. For each feature, the total weighted reduction across all the nodes represent its importance for corresponding decision tree. For an ensemble of trees, we can estimate the actual feature importance by averaging the feature importance from individual decision trees (Friedman 2001). We have used the feature score provided by XGBoost (Friedman et al. 2000) to assess the relative importance of used features.

### Performance Evaluation

The stakes associated with decision errors by classifiers are not always symmetric across possible class labels. Using our tool, later we have plan to build a repository of high quality indels enriched with true positives (TP). To achieve this, we have been more interested on the reduction of false positive (FP) calls than that of false negative (FN) calls. Thus, we have reported precision or positive predictive value (PPV) as our main performance metric which is defined as follows: *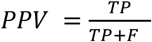*

For every sequencing pipeline of a particular platform, an aligner and a variant caller mentioned in ***Supplementary Table 1***, we have calculated a base precision, intersection (using RTG tools (Cleary et al. 2015)) between indel callset generated by the sequencing pipeline and the gold standard GIAB indel callset. We have applied our classifier in two different settings. First, the classifier is trained using the gold standard indel calls (all or only platform generated) from one sample and tested on the indel callset of another sample produced by that particular pipeline. Unless otherwise specified, this is the default settings. Second, we have also performed train-test split using the pipeline generated indel callsets for each sample to discard the effect of common instances between the training and test samples. This was only done (c.f., the analysis of ***Figure 2***) to understand the classifier performance under completely randomized and controlled scenario. But this is not likely to be an actual application scenario.

### Tuning Hyperparameters

We have performed a grid search in the hyperparameters’ space for the fine tuning of our classifier. Three most important hyperparameters are:

1. Number of trees: this is the number of boosting rounds.
2. Tree depth: the maximum number of conditions checking from the root to a leaf
3. Learning rates: shrinkage factor for a new tree

Number of trees or boosting rounds for our gradient boosting machine is one of the most important predefined hyperparameters. ***Supplementary Figure 2*** shows the effect of increasing number of trees over performance. Evidently, initial boosting rounds improve performance drastically whereas later rounds show performance saturation.

Other two significant hyperparameters are learning rate and tree depth. ***Supplementary Figure 3*** explores their mutual effect over performance. Increasing tree depth generally boosts performance. But similar to number of trees, the performance increment saturates with increasing tree depth. ***Supplementary Figure 3*** shows that influence of learning rates and tree depth on performance is intertwined. Low learning rates slow down the performance saturation. So, attaining a high-performance mark becomes possible. Higher learning rates tend to saturate performance quickly.

Chosen values for our experiments are as follows, boosting round = 200, tree depth = 16 and learning rate = 0.25.

## Conclusions

Our findings in this paper should help researchers to identify a more reliable set of indels. This study also sheds lights on the genomic contexts likely to be responsible for interesting mutation events. We have future plans to use our variant filter to extract more complicated class of variants, e.g., multi-allelic indels. This study should also help to elucidate mutation history.

## Supplemental Materials

**Supplementary Table 1:** List of sequencing platform and variant caller combinations used in our experiment.

| **Platform** | **Caller** |
| --- | --- |
| GIAB Consensus | Gold-standard |
| Complete Genomics | CGATools |
| 10X Genomics | GATK |
| Illumina Paired End | FreeBayes |
| Illumina Paired End | GATK |
| Illumina Mate Pair | FreeBayes |
| Illumina Mate Pair | GATK |
| Illumina Long Read | FreeBayes |
| Illumina Long Read | GATK |

**Supplementary Table 2:**
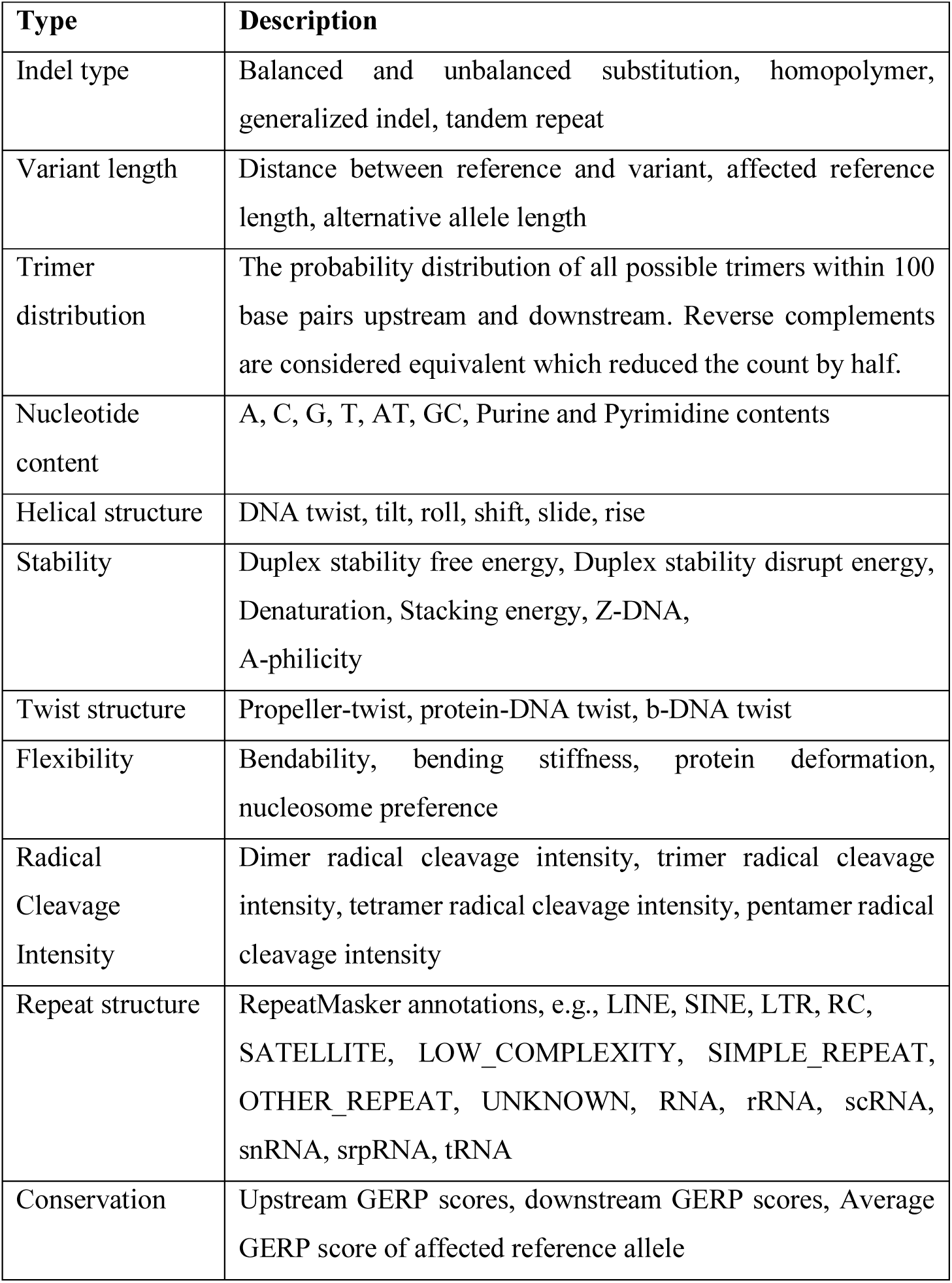
Selected features used for sample annotation.

**Supplementary Table 3:**
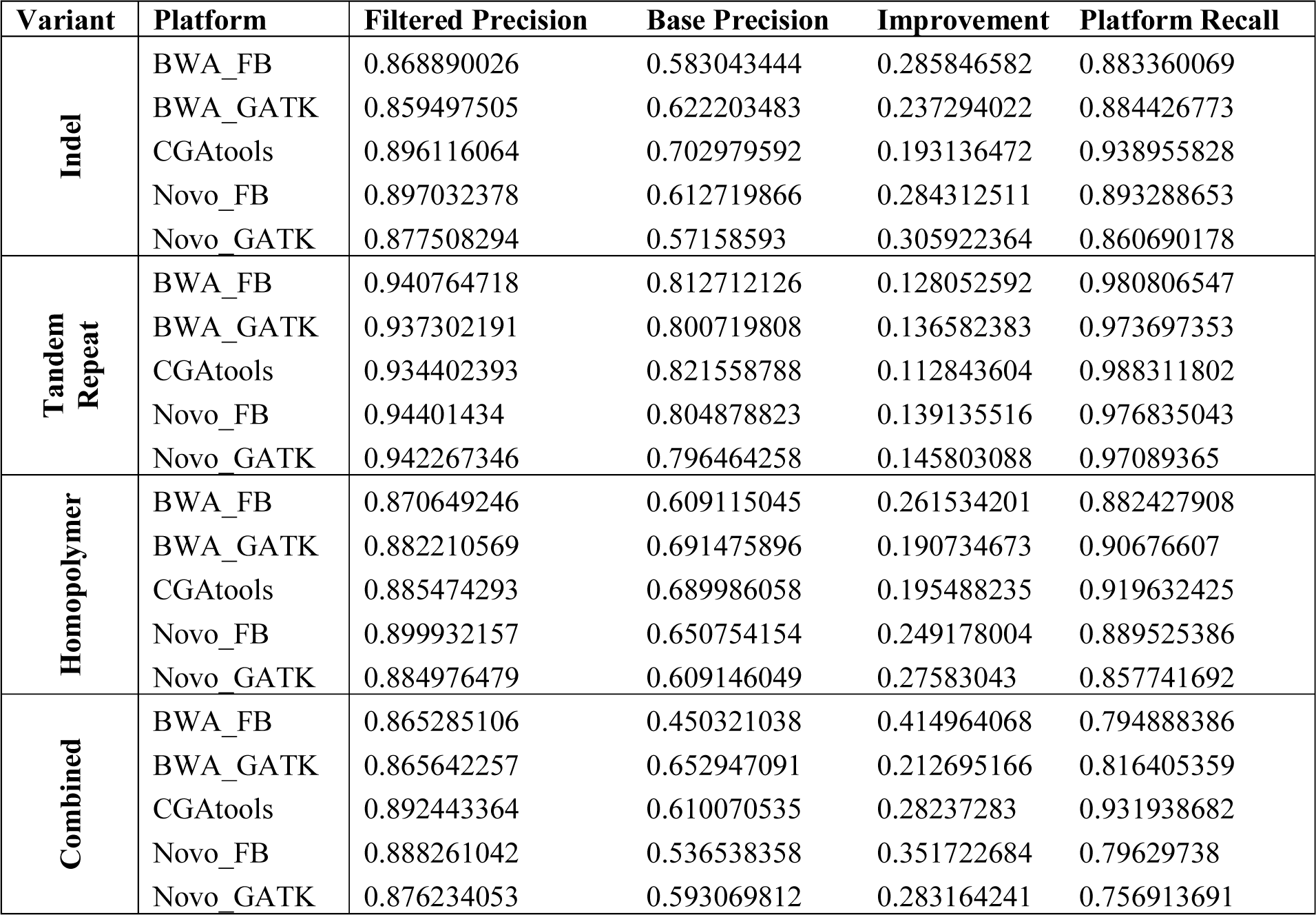
Precision (both before and after Filtering) and Recall across Different Platforms and Variant Categories (Training with HG003, HG004, HG005 and Testing with HG002)

**Supplementary Table 4:** Precision (both before and after Filtering) and Recall across Different Platforms (Training with HG001 and Testing with HG002)

| Platform | Filtered Precision | Base Precision | Improvement | Platform Recall |
| --- | --- | --- | --- | --- |
| 10x_Genomics | 0.829839207 | 0.654981899 | 0.174857308 | 0.882151456 |
| Complete_Genomics | 0.848867008 | 0.610070535 | 0.238796474 | 0.890191933 |
| DeepVariant_PrecisionFDA | 0.818655218 | 0.634213021 | 0.184442197 | 0.862526114 |
| IlluminaPairedEnd_FreeBayes | 0.838743642 | 0.530593095 | 0.308150546 | 0.878192968 |
| IlluminaPairedEnd_GATK | 0.820186934 | 0.557212315 | 0.262974618 | 0.846778272 |

**Supplementary Table 5:** Data for Figure 2 [Left panel].

| No of Samples | Platform-TP | GIAB-TP |
| --- | --- | --- |
| 0 | 0.610070534578925 | 0.610070534578925 |
| 1 | 0.849362313633914 | 0.827872729325533 |
| 2 | 0.87795531017638 | 0.850080457715 |
| 3 | 0.892443364103996 | 0.86390142852011 |
| 4 | 0.894835812876568 | 0.868815875200175 |

**Supplementary Table 6:** Data for Figure 2 [Right panel].

| Training Sample | Overlapping | Non-overlapping |
| --- | --- | --- |
| HG001 | 0.849266647064217 | 0.822827813535822 |
| HG003 | 0.867073369975678 | 0.836731004348079 |
| HG004 | 0.871835541071362 | 0.83580797576253 |
| HG005 | 0.849364939432526 | 0.825721263977402 |

**Supplementary Table 7:**
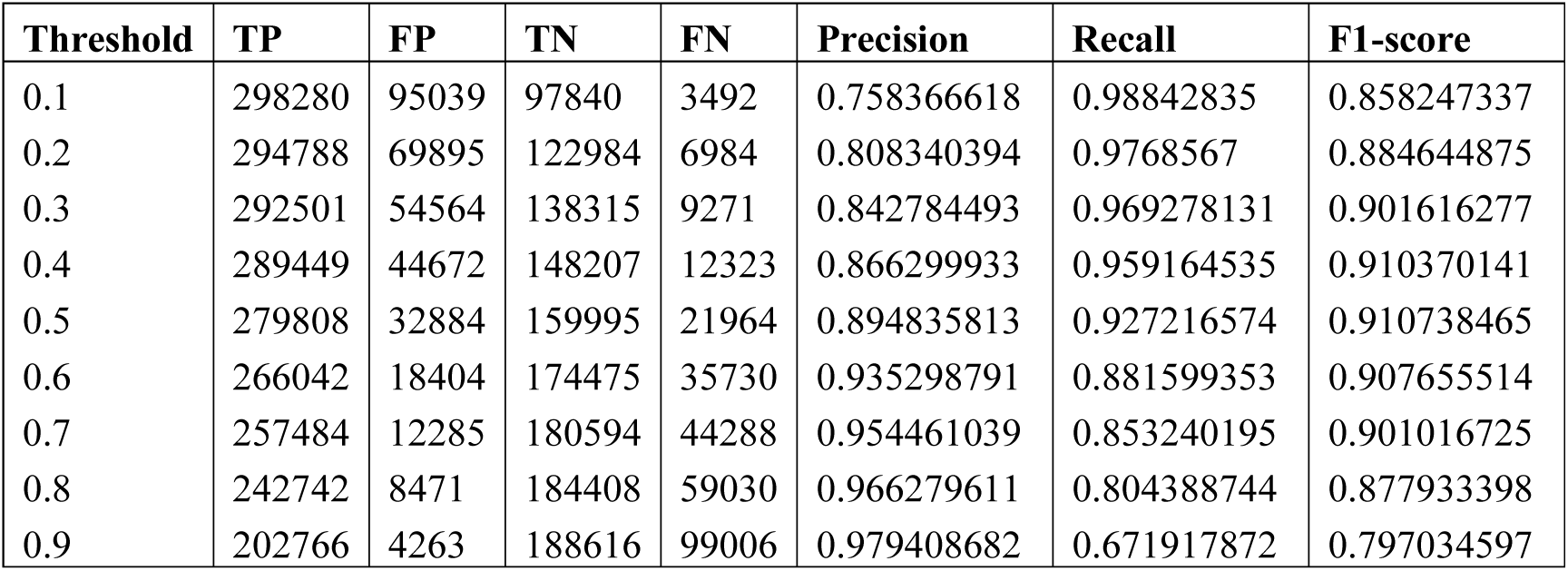
Data for Figure 3.

**Supplementary Table 8:** Data for Figure 5.

| Feature | Mean | STD |
| --- | --- | --- |
| GERP Score | 0.893857254 | 0.069916296 |
| DNA Structure | 0.687149784 | 0.112285741 |
| DNA Flexibility | 0.575843505 | 0.101948985 |
| ±100bp Composition | 0.524790685 | 0.106620855 |
| ±10/50bp Composition | 0.432841764 | 0.16987573 |
| DNA Stability | 0.392034154 | 0.073936636 |
| ALT Allele Composition | 0.089534154 | 0.083501268 |
| REF Allele Composition | 0.082080024 | 0.057725063 |
| ±5bp Composition | 0.036398041 | 0.012786659 |
| Repeat Structure | 0.026875692 | 0.032858894 |

**Supplementary Table 9:**
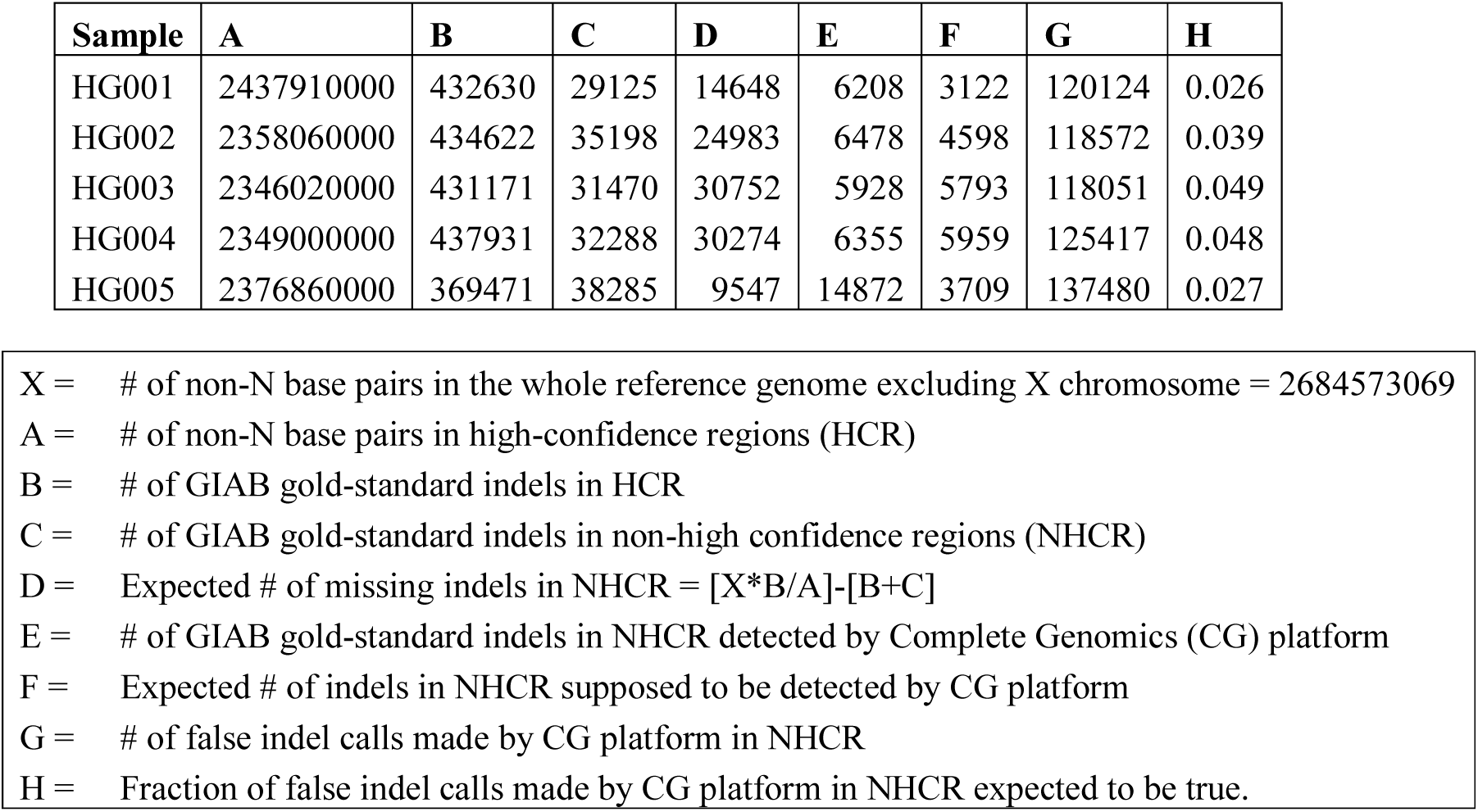
Calculation of the Fraction of False Indel Calls Made by Complete Genomics Platform in non-High Confidence Regions Expected to be True Figure 3.

| Sample | A | B | C | D | E | F | G | H |
| --- | --- | --- | --- | --- | --- | --- | --- | --- |
| HG001 | 2437910000 | 432630 | 29125 | 14648 | 6208 | 3122 | 120124 | 0.026 |
| HG002 | 2358060000 | 434622 | 35198 | 24983 | 6478 | 4598 | 118572 | 0.039 |
| HG003 | 2346020000 | 431171 | 31470 | 30752 | 5928 | 5793 | 118051 | 0.049 |
| HG004 | 2349000000 | 437931 | 32288 | 30274 | 6355 | 5959 | 125417 | 0.048 |
| HG005 | 2376860000 | 369471 | 38285 | 9547 | 14872 | 3709 | 137480 | 0.027 |

|  |  |
| --- | --- |
| X = | # of non-N base pairs in the whole reference genome excluding X chromosome = 2684573069 |
| A = | # of non-N base pairs in high-confidence regions (HCR) |
| B = | # of GIAB gold-standard indels in HCR |
| C = | # of GIAB gold-standard indels in non-high confidence regions (NHCR) |
| D = | Expected # of missing indels in NHCR = $[X*B/A]-[B+C]$ |
| E = | # of GIAB gold-standard indels in NHCR detected by Complete Genomics (CG) platform |
| F = | Expected # of indels in NHCR supposed to be detected by CG platform |
| G = | # of false indel calls made by CG platform in NHCR |
| H = | Fraction of false indel calls made by CG platform in NHCR expected to be true. |

**Supplementary Figure 1:**
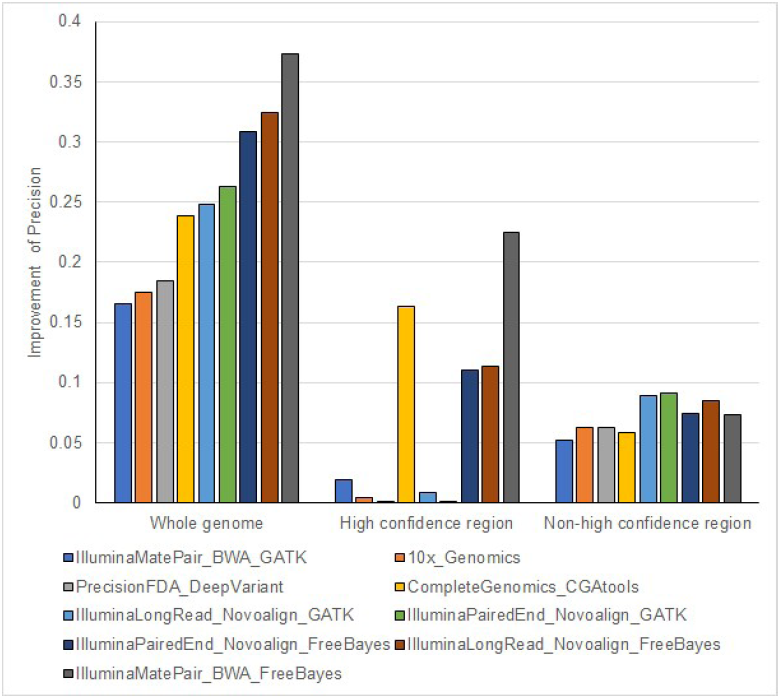
Precision Improvement over both Whole Genome and High/non-High Confidence Regions Separately for Different Platforms.

**Supplementary Figure 2:**
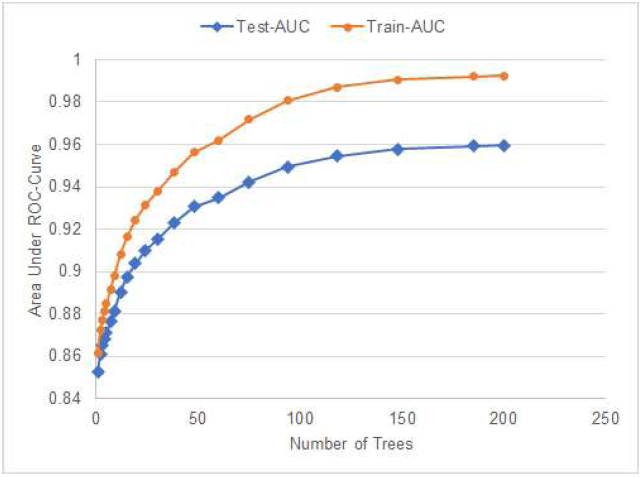
Area under ROC curve vs. Number of Boosting Trees.

**Supplementary Figure 3:**
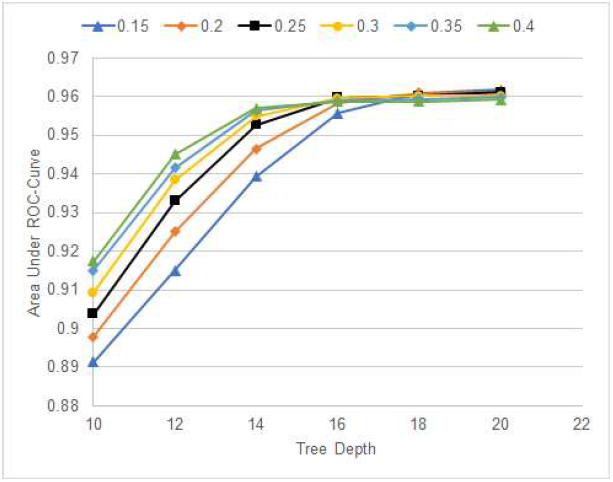
Area under ROC curve vs. Tree Depth for Different Learning Rates.

**Supplementary Figure 4:**
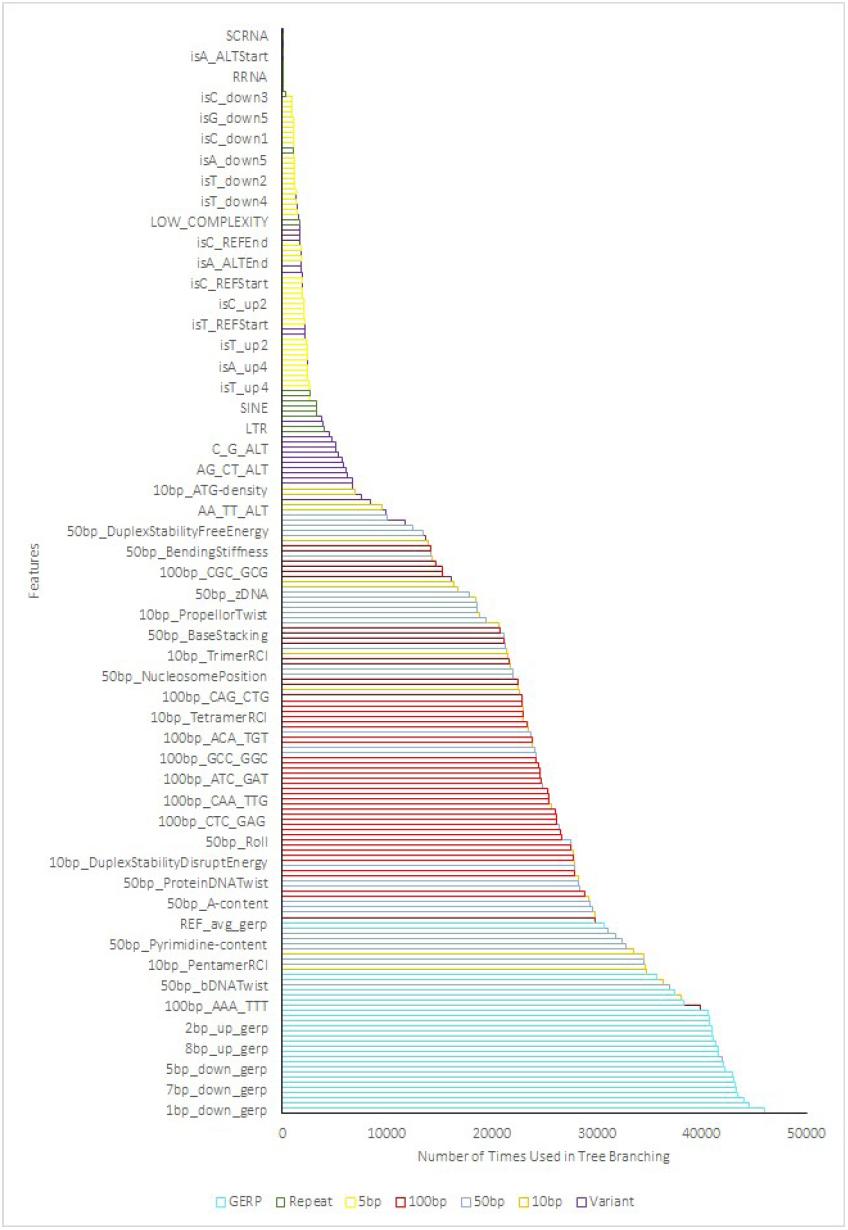
Usage Count of Classifier Features in Sorted Order.

1 https://precision.fda.gov/challenges/truth/results

2 https://www.illumina.com/science/technology/next-generation-sequencing/mate-pair-sequencing.html

3 https://www.illumina.com/science/technology/next-generation-sequencing/long-read-sequencing.html

4 http://www.novocraft.com/products/novoalign/

5 https://precision.fda.gov/challenges/truth/results

